# Comparative analyses of 3654 chloroplast genomes unraveled new insights into the evolutionary mechanism of green plants

**DOI:** 10.1101/655241

**Authors:** Ting Yang, Xuezhu Liao, Lingxiao Yang, Yang Liu, Weixue Mu, Sunil Kumar Sahu, Xin Liu, Mikael Lenz Strube, Bojian Zhong, Huan Liu

## Abstract

**Background:** Chloroplast are believed to arise from a cyanobacterium through endosymbiosis and they played vital roles in photosynthesis, oxygen release and metabolites synthesis for the plant. With the advent of next-generation sequencing technologies, until December 2018, about 3,654 complete chloroplast genome sequences have been made available. It is possible to compare the chloroplast genome structure to elucidate the evolutionary history of the green plants.

**Results:** We compared the 3654 chloroplast genomes of the green plants and found extreme conservation of gene orders and gene blocks in the green plant such as ATP synthase cluster, Phytosystem, Cytochrome cluster, and Ribosomal cluster. For the chloroplast-based phylogenomics, we used three different data sets to recover the relationships within green plants which accounted for biased GC content and could mitigate the bias in molecular data sets by increasing taxon sampling. The main topology results include: I) Chlorokybales + Mesostigmatales as the earliest-branching lineage and a clade comprising Zygnematales+ Desmidiales formed a grade as the sister group to the land plants, II) Based on matrix AA data, Bryophytes was strongly supported as monophyletic but for matrix nt123 data, hornworts, mosses and liverworts were placed as successive sister lineages of *Tracheophytes* with strong support, III) Magnoliids were placed in the outside of Monocots using the matrix nt123 data and the matrix AA data, IV) Ceratophyllales + Chloranthales as sister to the Eudicots using matrix nt123 data, but when using matrix nt12 data and AA data, only Ceratophyllales sister to the Eudicots.

**Conclusion:** We present the first of its kind large scale comparative analyses of the chloroplast coding gene constitution for 3654 green plants. Some important genes likely showed co-occurrence and formed gene cluster and gene blocks in Streptophyta. We found a clear expansion of IRs (Inverted Repeats) among seed plants. The comprehensive taxon sampling and different data sets recovered a strong relationship for green plants.

## Introduction

Chloroplast are one of the most important organelle of land plants and green algae which are essential for plant growth and development, and are involved in photosynthesis, lipid metabolism, and other cellular processes. According to endosymbiotic theory, chloroplast originated from a cyanobacterial endosymbiont into a eukaryotic host, and the photosynthetic eukaryotes were endosymbionts of non-photosynthetic eukaryote hosts to form secondary chloroplast, then the cellular chimaera subsequently diversified into glaucophytes, red algae, and green plant/algae. However, it is unclear whether the chloroplast of red algae, green algae, and green plants were from a single origin or multiple origins [1]. As the transfer of the chloroplast genes to the nucleus was an ongoing process, the phylogenetic tree based on some chloroplast genes may be complex, if sequences involved in the analyses are from different origins. However, chloroplast genomic DNA (cpDNA) are conserved in gene content, the similar set of genes in cpDNA could be explained in terms of large-scale gene transfer in an ancestral lineage and could help us to understand chloroplast origin and evolution. For instance, the presence of gene clusters like psbB/T/N/H could be considered as an indication of monophyly [2, 3].

cpDNA of green plants (Viridiplantae) normally exhibit a conserved genome structure which contains two copies of an inverted repeat (IR) separating the small single-copy region (SSC) and the large single-copy region (LSC). The chloroplast genome size of green plants normally ranges from 107 kb (*Cathaya argyrophylla*, Pinaceae family) [4] to 218_L_kb (Pelargonium, Geraniaceae family) [5]. However, some angiosperm lineages may have extreme variations in their genome size, for instance, *Cutinus* (Cytinaceae) chloroplast genome is around 20 kb, while Chlorophyta (i.e., *Floydiella*, Chaetopeltidaceae) chloroplast genomes have been reported to host an unexpected large size of 520 kb [6]. The size of cpDNA has been compared within many clades [7, 8], and many factors could explain these chloroplast genome size variation, like (a) variations of intergenic regions, intron lengths, etc. [9, 10]; (b) IR region variation [5, 11]; (c) gene loss [12].

For cpDNA of green plants, hotspots for structural variation include the IRs, gene loss, gene transfer, and gene arrangement. For the IR variation, the lengths of IRs are likely to be expanded, contracted or to be completely lost. The IR analyses of all green plants showed that short IRs are frequently found in Bryophyta followed by Chlorophyta, the lowest among Polypodiopsida followed by basal Magnoliophyta, Magnoliidae, Commelinids [11], and in Papilionoideae, Pinaceae, cupressophytes, IRs are nearly lost or missing [8, 13, 14]. Regarding the gene variation, the cpDNA of green plants are normally conserved, but gene losses are widely seen especially in parasitic plants such as *Cuscuta* and *Epifagus*, which have partially or completely lost the photosynthetic ability [15].

To understand the origin and relationships of green plants, the phylogenetic analyses have been widely performed based on nuclear, mitochondrial [16], and chloroplast loci [17, 18]. The phylogenetic relationship among Chlorophyta has been reviewed recently [19–22] and the branching orders of the prasinophyte lineages, the relationships among core chlorophyte clades (Chlorodendrophyceae, Ulvophyceae, Trebouxiophyceae and Chlorophyceae) required further deep analyses. Meanwhile, regarding the ferns [23, 24], and Bryophytes [25, 26], transcriptome sequencing data was used to resolve the debated topologies within the ferns and Bryophytes. For the gymnosperm group, Lu et al. (2014) used two nuclear genes and performed near complete sampling of extant gymnosperms genera, and found that the cycads are the basal-most lineage of gymnosperms rather than a sister to Ginkgoaceae, a sister relationship between Podocarpaceae and Araucariaceae [27]. For seed plants, Burleigh et al. used four nuclear loci, five chloroplast loci and four mitochondrial loci from 31 genera to resolve the seed plant tree of life [17]. For basal angiosperms, Moore et al. used 61 chloroplast genes from 45 taxa to reconstruct the phylogenetic order among basal angiosperms [28]. Likewise, the largest chloroplast phylogenetic study has been performed across green plants by using a nearly complete set of protein-coding sequences, based on 360 species of the green plants, and 1879 taxa representing all the major subclades [29, 30].

With the advent of next-generation sequencing technologies, enormous efforts have been made to sequence the whole chloroplast genomes of plants. Until December 2018, over 3,000 complete chloroplast genome sequences have been made available in the National Center for Biotechnology Information (NCBI) organelle genome database. This large amount of complete cpDNA sequences could be effectively utilized to understand the evolution of the chloroplast genomes and phylogenetic relationships among plants. With so many chloroplast genomes, we tried to answer three main questions from this study: i) After the split of Streptophyta and Chlorophyta, how the evolution shaped Streptophyta and what were the similarities the in the genome? ii) IRs degenerated widely in red algal and have uneven size distribution in Viridiplantae, what is the formation mechanism behind IRs? iii) does increasing taxon sampling would help to resolve phylogenetic questions of relationships in Viridiplantae? In the present study, we comprehensively analyzed the available chloroplast genomes of Viridiplantae comprising 3,654 taxa, 298 families, and 111 orders. We compared the genomic organizations in their cpDNAs between major clades, including gene gain/loss, gene copy number, GC content, gene cluster, and gene blocks. We also covered a wide range of green plants species to construct the chloroplast-based phylogenetic trees. Increasing taxon sampling together with the whole coding genes of chloroplast helped us to resolve phylogenetic questions of relationships in Viridiplantae.

## Results

### General Characteristics of the Genomes

#### The genome size and gene organization in chloroplast genomes

In this study, the complete chloroplast genomes (cpDNA) of 3,654 taxa representing 298 families, and 111 orders were selected. The size of cpDNA ranged from 71,666 bp to 521,168 bp. Liverworts, mosses, and gymnosperm displayed the smallest average genome size, while Chlorophyta had the largest genome size variation. Even though the chloroplast genome size showed large variation, but there were 120–130 conserved genes as well. We recovered 79 protein-coding genes from all the sequenced cpDNA, but seven genes: *ndhF*, *psaA*, *psaB*, *rpoB*, *rpoC1*, *rpoC2*, *ycf2* had no information regarding gene annotation (Liu et al 2019) (see methods), so we only investigated 72 protein-coding genes in the follow-up analysis.

To investigate the gene content similarity in the green plants, we calculated the average gene number in every order and the overview of the genes are presented in Fig 1. According to the copy number of the gene, samples were divided into three main clades, the first clade comprised some of Chlorophyte, Charophyta, moss, liverworts, fern-and-horsetails, and the second contained most of Chlorophyta, Genetal and Pinales of Gymnosperm, Santalales of Eudicots which possessed no ndh family. The third is Eudicots and Monocots which contained two copies of *rpl23*, *rpl2*, *ndhB*, *rps7*, and *rps12*.

**Fig 1.**
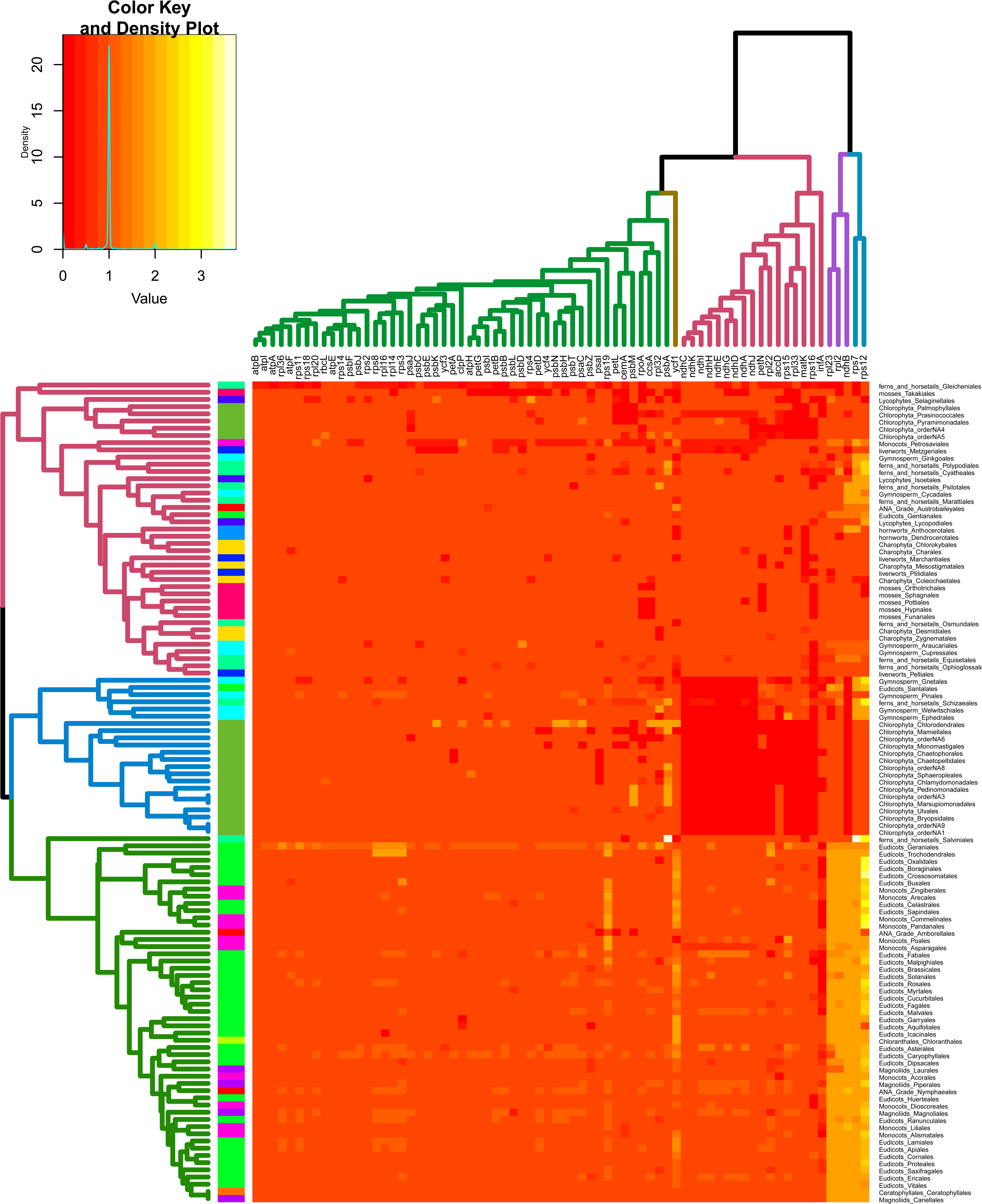
The gene constitution in the green plants is displayed as a heatmap. In heat map, the data is displayed in a grid where each row represents order and each column represent average gene number in the order.

We also compared the gene content in the green plants by using the Spearman correlation (Fig S1). Some genes appeared to co-occur, the NADH dehydrogenase (ndh) genes showed a strong positive correlation with other genes within the family (r>0.7) except ndhB (r ∼0.5), however, ndhB showed a strong positive correlation with *rpl2*, *rpl23*, *rps12*, *rps7*, and *ycf1*. Ndh family showed a negative correlation with both *clpP* and *infA* genes. We also found that some families were closely correlated such as psb family (*psbE*, *psbF*, and *psbH*) had a positive correlation to pet family (*petA*, *petB*, *petD*, *petG*, and *petL*).

Similarly, the introns in land-plant chloroplast genomes are generally conserved. The Chlorophyta and Charophyta possess the least intron number. Most of the genes lacked introns with the exception among several ribosomal proteins and photosynthesis genes such as *atpF*, *ndhA*, *petD*, *rpl16* and *rps12* which possess at least one intron (in Streptophyta). The intron number of *clpP* gene showed a high divergence, 2327 species showed the presence of two introns, and more than 100 species owned 3-4 introns. Moreover, most of the *clpP* gene without introns were found among Chlorophyta, gymnosperms and Poaceae (Fig S2, Table S1).

#### Gene gain/loss in chloroplast genomes

Although the genetic content and number of protein-coding genes are generally conserved in the chloroplast genomes, the gene gain and loss have been reported. A total of 72 protein-coding genes were investigated from 3654 species (Table S1).

For gene gain, we found that, from Nymphaeales, almost all the flowering plants have two copies of ndhB, rpl2, rpl12, rpl23, rps12, and rps7 genes which correlated with IRs expansion, especially rps12 with four copes. In Campanulaceae, Ericaceae and Fabaceae, ndh family genes were duplicated.

For gene loss, we found some genes were more likely lost in the green plant. The chloroplast translation initiation factor 1 (*infA*) and the ribosomal protein L22 (*rpl22*) are the two housekeeping genes. We found that *infA* was absent in 1825 taxa, and it was more frequently observed among angiosperms, especially in Eudicots. On the other hand, *rpl22* was missing in 474 taxa mainly in Chlorophyta and legumes, suggesting that both genes were possibly transferred from chloroplast to the nucleus during evolution, as reported in an earlier study [31]. The ndh genes are related to the cyclic electron in the photosystem I complex, and has been thought to be lost in the higher plants [32]. In our study, ndh genes were found to be lost in at least 300 species, mainly in Chlorophyta, Pinaceae, Ephedrales, Welwitschiales, Gnetales and some species of Orchidaceae. At the same time, except ndh gene family, *petN*, *rpl22*, *rpl33*, *rps15*, *rps16* were lost in Chlorophyta, and *rps16* were lost in Gymnosperm and Bryophytes. In addition, we also observed the loss of *accD* and *ycf1* genes together in more than 800 species (almost are Poales). *Ycf1* is thought to have a functional role of assembling the ACCase holoenzyme [33, 34], and both these two genes have been related to fatty acid synthesis.

#### Gene conservation and rearrangement

It is well known that the structure of chloroplast genomes conserved and the order of genes is relatively consistent in land plants. This opens up the possibility of reconstructing insertions, deletions and inversions during the evolution of green plant. In this study, 72 protein-coding genes were ordered according to the annotated position.

In *Arabidopsis thaliana*, cluster analysis has been done based on chloroplast transcriptomes expression and finally chloroplast genes divided into eight sub-clusters [35]. To calculate the blocks frequency in Streptophyta, we first removed the samples in the order which have the same gene content, and finally obtained 1517 cpDNA and the blocks frequency are listed in Table S2. Based on the functional categories, there are three major gene clusters. The frequency of ATP synthase cluster: *atpA-atpF-atpH-atpI* was 74%, *atpE-atpB* was 82%, Phytosystem and Cytochrome cluster: *petA-psbJ-psbL-psbF-psbE-petL-petG* was 80% *psbB-psbT-psbN-psbH-petB-petD* was 85%. Ribosomal cluster: *rps8-rpl14-rpl16-rps3* was 83%, *rpl33-rps18-rpl20* was82% and *rpoA_rps11_rpl36* was 85%. In Monocots and Eudicots, we observed three photosystem gene clusters with high frequency: *psbM/D/C/Z* [60%], *psbJ/L/F/E* [85%] and *psbB/T/N/H* [88%]. *PsbJ/L/F/E* and *psbB/T/N/H* nearly conserved in all the green plants and liked to form blocks: *psbB/T/N/H-petB-petD-rpoA-rps11-rpl36* [78%], *psbJ/L/F/E-petL-petG-psaJ-rpl33-rps18-rpl20* [76%] in Streptophyta.

But *psbM/D/C/Z* block showed the highest variability in the green plant. *PsbD* and *psbC* genes encode the D2 and CP43 proteins of the photosystem II complex, and they are generally co-transcribed [36]. Similarly, *psbM* is highly light-sensitive and plays important roles in such conditions, in fact, the knock-out of *psbM* leads to a significant decrease in the activity of photosystem II [37]. In Chlorophyta, *psbD/C/Z*, *psbZ/M*, and *psbD/C* were found to be widely distributed, but in the Charophyta branch, only *psbD/C/Z* block exists. Later in Bryophytes, *psbZ/C/D* and *psbM* were connected by ATP synthase: *atpA/F/H/I*. From ferns and horsetails clade, *psbM/D/C/Z* was formed. In Cycadales which likely represents all the seed plant, showed the presence of complete *psbM/D/C/Z* blocks, but in Pinales, *psbM* and *psbD/C/Z* were separated. In Poaceae *atpA/F/H/I-rps2-petN-psbM* was especially inverted leading to the production of larger block *psbK/I/M/D/C/Z*.

Except gene cluster, there are still some large blocks which contained the more than one functional category gene. The largest block: (*atpA-atpF-atpH-atpI*) (1) - (*rps2-petN-psbM*) (2) -(*psbD-psbC-psbZ*) (3) -(*rps14-ycf3-rps4*) (4) [51%]- (*ndbJ-ndhK-ndhC-atpE-atpB-rbcL*) (5) [70%]*-accD-psaI-(ycf4-cemA-petA-psbJ-psbL-psbF-psbE-petL-petG-psaJ-rpl33-rps18-rpl20*) [69%]- (*psbB-psbT-psbN-psbH-petB-petD-rpoA-rps11-rpl36*) [78%] was found with high frequency in Streptophyta, and numbers in [] are the block frequency. In Poaceae, (1), (2) and (3) block were with complicated situation which experienced several rounds of recombination and in some genus of Fabaceae, (3) -(4) and (5) were cross interchanged. In streptophytes, the cpDNA reserved a large gene block:(*psbB-psbT-psbN-psbH-petB-petD*) [85%]-(*rpoA-rps11-rpl36*) [85%]-*infA*-(*rps8-rpl14-rpl16-rps3-rpl22-rps19-rps2-rps23*) [61%] which included S10-spc regions and connected directly with IRb (Table S3).

IRs normally contain tRNAs and rRNAs, but in this study, we didn’t annotate tRNA and rRNAs, we focused on coding genes: *rps19-rpl2-rpl23-ndhB-rps7-rps12* which were newly acquired in IRs for angiosperms [8] (Fig 2). The *rps19-rpl2-rpl23* were conserved in the green plants, but *ndhB-rps7-rps12* showed great variation. The *ndbB* were found in some green algae such as Prasinococcales and Palmophyllales, then from Charophyta, *ndhB-rps7-rps12* and *rps19-rpl2-rpl23-ndhB-rps7* blocks were formed. In the Bryophytes and Lycophytes, *ndhB-rps7-rps12* widely existed and Dendrocerotaceae have a complete block of *rps19-rpl2-rpl23-ndhB-rps7-rps12*. In ferns and horsetails, except *rps19-rpl2-rpl23-ndhB-rps7-rps12* block in Marattiales, most orders have *ndhB-rps12-rps7-psbA-ycf1* block which is near the IRs regions. In Amborella and Nymphaeales, IRs contained *rps19-rps2-rps23-ndhB-rps7-rps12* which is similar to all other angiosperms but *Trithuria filamentosa* and *Cabomba caroliniana* had the largest IRs which contained *psbB-psbT-psbN-psbH-petB-petD-rpoA-rps11-rpl36-infA-rps8-rpl14-rpl16-rps3-rpl22-rps19-rps2-rps23-ndhB-rps7-rps12*.

**Fig 2.**
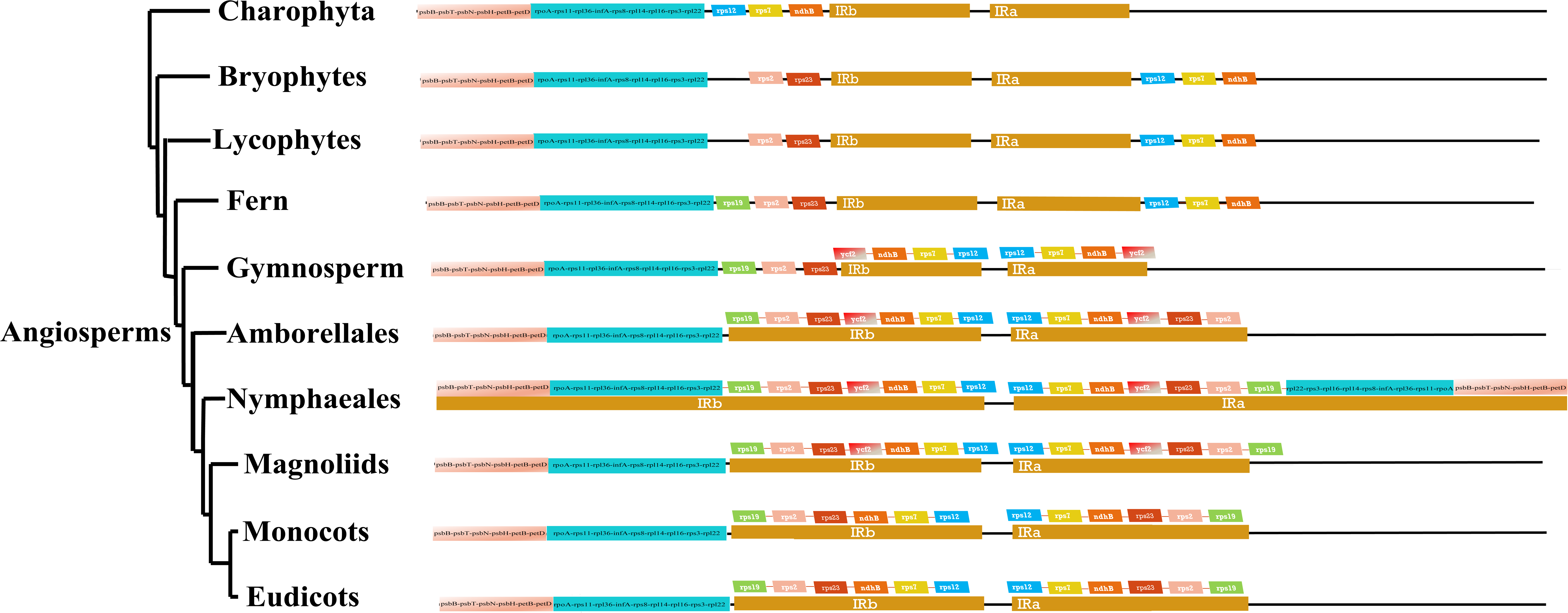
Coding genes in IRs in Streptophyta. Coding genes in IRs and upstream are shown.

#### GC bias

The GC content is deemed to be in connection with the amino acid composition based on former research[29]. In this study, the GC content at different codon positions of all the 72 protein-coding genes including the first, second, and third codon positions, together with ntNo3rd and ntAll data sets were used for the analysis. The average GC content of ntNo3rd matrix ranged from 36.3% to 58.2%, while the ntAll data set had a slightly lower GC content of 38.2%, varying from 29.0% to 55.4% (Table S1). The third codon position of all 14 clades owned the lowest GC content compared to other position, and there was an obvious difference between different clades. Therefore, the One-Way ANOVA was carried out using SPSS to test whether there was any significance between different clades, and the result revealed that the GC characters were the same in Chlorophyta and Charophyta, and there was a non-significant difference between angiosperms. Furthermore, the GC content of Lycophytes and fern were significantly higher than all the other clades (p < 0.01), and the GC content in the clades after Lycophytes and fern were higher than that of Bryophytes and algae (Fig S3).

#### Phylogenetic analysis

To conduct the phylogenetic analysis, the concatenated alignment of three data sets for the 72 genes from 3654 species were used with 6 Rhodophyta as outgroups, which consisted of 4,724 amino acid positions (AA). A total of 44,187 positions for the matrix containing all codon positions (nt123) and 29,458 positions for the matrix containing all but the third codon positions (nt12). We used two programs: IQ-TREE and RAxML to construct the phylogenetic tree, but they both produced nearly the same topology, so we only used IQ-TREE to illustrate our results.

The topology is summarized in Fig 3-4 and the details of the phylogenetic trees are provided in supplemental materials (Fig S4-S8). For some debated clades, the summary of the similarities and conflicts in topologies derived from these three data sets are presented in Table 1. All phyla of green plants except Charophyta are recovered as monophyletic. Within Chlorophyte, both matrix nt12, nt123 and the matrix AA supported that Palmophyllales and Prasinococcales are both the earliest-diverging lineage of the Chlorophyta (SH-alrt == 33%, UFboot = 100%). Chlorophyceae is strongly supported as monophyletic, with Chlamydomonadales + Sphaeropleales sister to a clade of Chaetophorales and Chaetopeltidales. The relationship of Chlorophyceae and Ulvophyceae (non-monophyletic) is uncertain.

**Fig 3.**
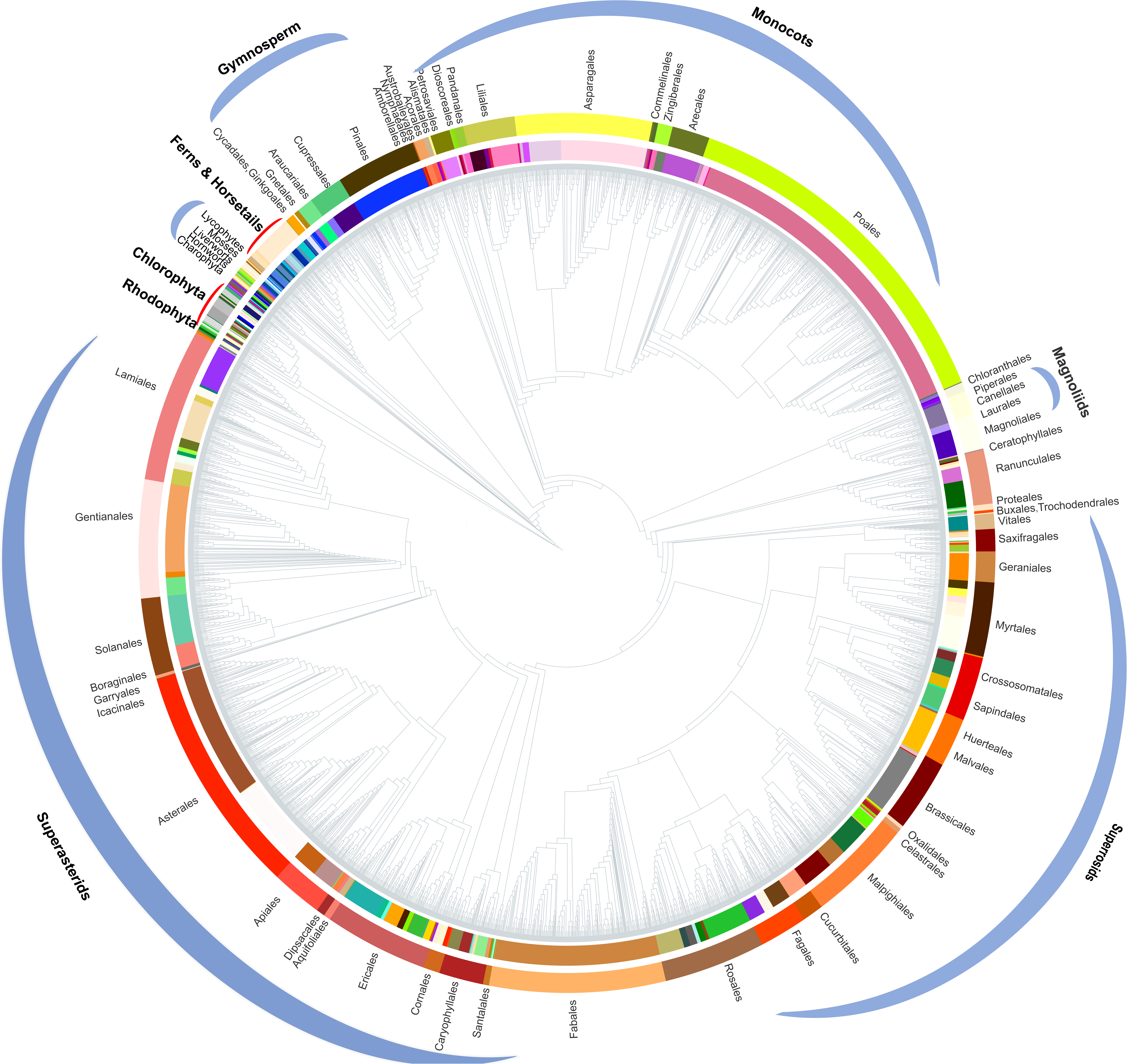
Chloroplast phylogenomic tree based on the matrix nt12 of 72 protein-coding genes of 3654 green plants and six Rhodophyta using IQTREE. The colors in the internal circle indicate different families while the colors in the external circle indicate different orders. The green branches represent the branch with UFboot more than 95%.

**Fig 4.**
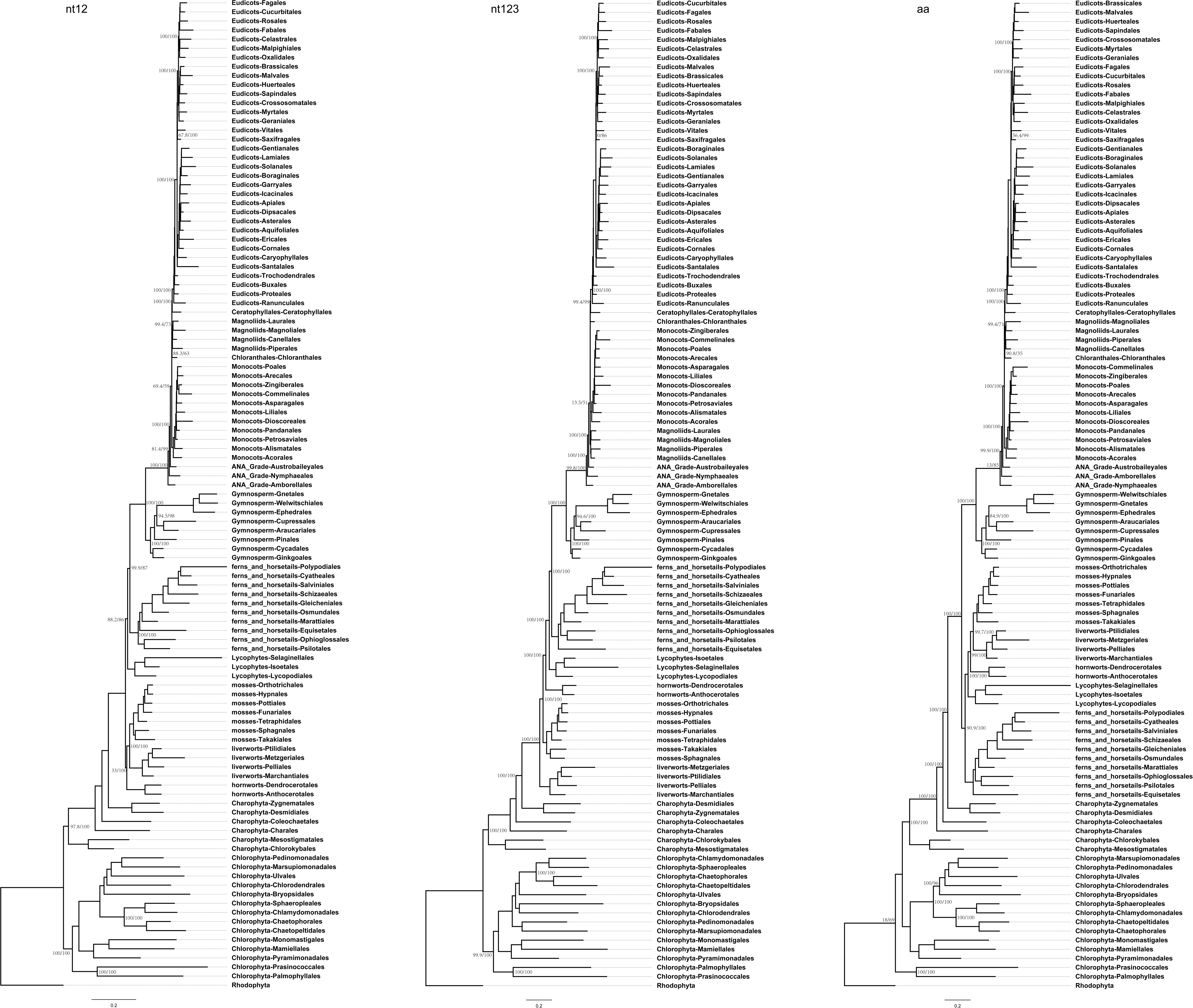
Summary of the phylogenomic tree based on three data sets of 72 protein-coding genes of 3654 green plants and six Rhodophyta using IQTREE.

Within Streptophyta, Charophyta lineages formed a paraphyletic assemblage with the land plants. Among the Charophyta groups, Chlorokybales + Mesostigmatales are the earliest-branching lineage and a clade of Zygnematales+ Desmidiales formed a grade as the sister group to the land plants.

Within land plants, Bryophytes were weakly supported as monophyletic based on matrix nt12 (SH-alrt == 33%, UFboot = 100%), while matrix AA, strongly supported Bryophytes as monophyletic (SH-alrt == 100%, UFboot = 100%), both matrix nt12 and matrix AA supported liverworts and mosses are uniting and sister to hornworts. For matrix nt123 data, hornworts, mosses and liverworts were found to be successive sister lineages of *Tracheophytes* (SH-alrt == 100%, UFboot = 100%).

Within *Euphyllophyta* in the matrix nt12 and nt123, a well-supported *Monilophyta* is sister to *Spermatophyta* (UFboot = 100%), but the matrix aa indicated that *Monilophyta* is sister to Bryophytes (SH-alrt = 90.9%, UFboot = 100%). Within *Monilophyta*, matrix nt12 supported Ophioglossales as the earliest-diverging lineage (UFboot =100%), but matrix nt123 supported Equisetales as the earliest branch (UFboot =100%). Within *Spermatophyta*, gymnosperms were designated as sister to angiosperms. Within gymnosperms, the subclades were well supported in three data sets, the clade of Cycadales + Ginkgoales is sister to the rest of gymnosperms. While the Gnetales, Welwitschiales along with Ephedrales, formed a clade (UFboot = 100%), which are sister to the clade comprising Cupressales and Araucariales.

Within angiosperms, in matrix nt12 and nt123, the Amborellales is recovered as the sister to all other angiosperms, followed by Nymphaeales, nevertheless, the placement of Nymphaeales is outermost in angiosperms in the matrix AA with weakly support (SH-alrt =34.9%, UFboot = 100%). The placement of Ceratophyllales is certain in the outside of Eudicots in the three data sets, while the Magnoliids is placed in the outside of monocots in matrix nt123 (SH-alrt =100%, UFboot = 100%), and nested between monophyletic monocot and eudicot in matrix nt12 (SH-alrt =64.9%, UFboot = 59%) and matrix AA (SH-alrt =90.8%, UFboot = 35%). While checking the amino acid composition in Magnoliids, we found the composition deviated significantly from the ‘master’ distribution using the chi^2^ test. The relationship between COM clade supported Oxalidales is sister to Celastrales + Malpighiales. The major subclades were typically well supported in Monocots and Eudicots, but the position of Vitales, Gentianales, Petrosaviales and Arecales remained problematic. To further verify the phylogenetic analysis, the data of amino acids from the former research were added into the tree construction [30], and the results showed that the data of the same orders are clustered together, and the topology of the major clade is consistent with the matrix nt12 (Fig S9).

#### Selective pressure of chloroplast genes

By dissecting the chloroplast genomes (cpDNA), we learned the special gene gain/loss, gene copy number, gene cluster order, and GC content along the evolution of the green plants, we also tried to understood which clades and cpDNA genes were under purifying selection, nonsynonymous substitution (dN), synonymous substitution (dS) and the dN/dS ratio were also calculated. The ndh subunits were thought to be related to the synthesis of photosystem I complex and involved in the adjustment of the redox level of the cyclic photosynthetic electron transporters [38]. Almost all the photosynthetic land plants have the ndh genes except in Chlorophyta, gymnosperms and some species of Orchidaceae (Table S1). To investigate the selective pressure of photosynthesis-related genes influenced by the absence of ndh subunits, 37 species in all 14 clades (10 of 37 species lost all ndh genes) were selected and the sequence of 14 photosynthesis genes existing in all the 37 species were chosen (Fig 5, Table S3). The results of the dN/dS analysis of 37 species revealed that the species which lost ndh genes might have a higher dN and dS, especially in Eudicots and gymnosperms. The dN/dS was extremely high in Chlorophyta, implying the occurrence of positive selection and dN/dS of all Streptophyta species except *Downingia cuspidata* is less than 1, suggesting that chloroplast genes are under purifying selection, even the rate was higher in the species without ndh genes. In addition, the independent analysis of 14 photosynthesis genes suggested that dS and dN in all genes for angiosperms have a relatively higher dN without ndh genes, except *psbK* (Fig S10, Fig S11).

**Fig 5.**
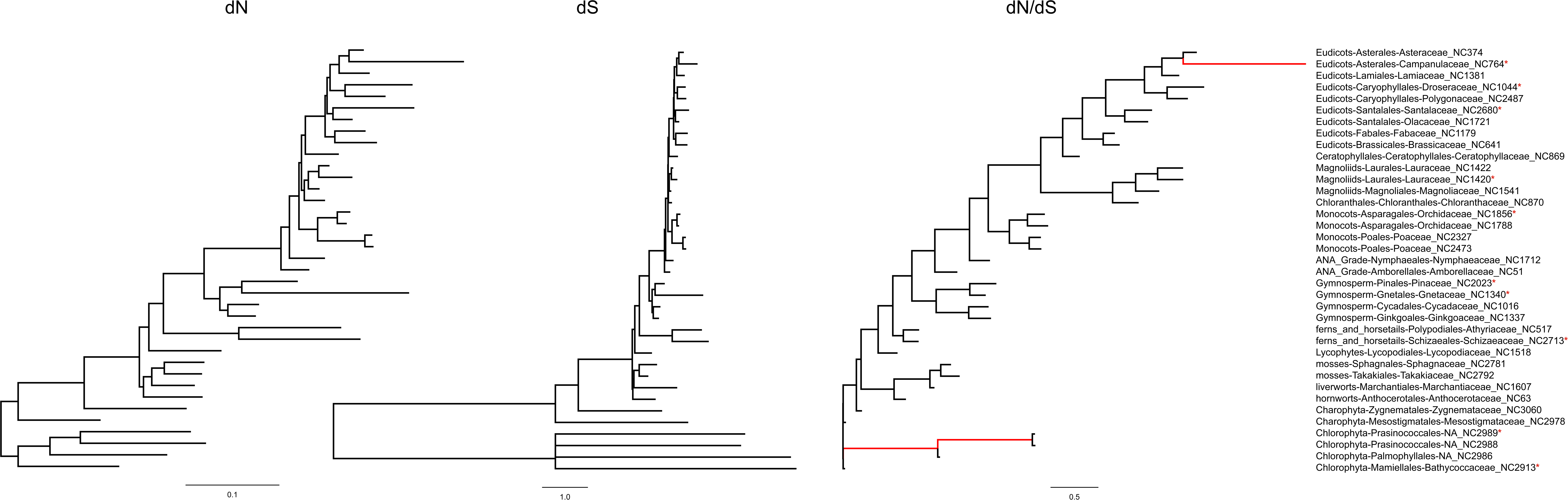
Phylogenomic tree of selective pressure for 37 species in all 14 clades. The red branch stands for the length of the branch which was changed to 1. “*” indicated the species without ndh genes.

## Discussion

### The green plant’s chloroplast genome

The chloroplast gene content in the green plants are conserved and plasmid genome architectures have been discussed based on recombination events of rDNA operons [39], but how similarity of the genome in Streptophyta is unknown. In our study, we found the same function class likely to form gene cluster, ATP synthase, Phytosystem and Cytochrome, Ribosomal cluster appeared more than one with high frequency. In Streptophyta, a block: (*psbB-psbT-psbN-psbH-petB-petD*) [85%] -(*rpoA-rps11-rpl36*) [85%]-*infA*-(*rps8-rpl14-rpl16-rps3-rpl22-rps19-rps2-rps23*) [61%] widely existed and was located nearly IRs regions, parts of them are the *S10–spc*–alpha operon locus which first appeared in eubacteria and archaebacteria. Euglena and glaucophyte plasmids in the *S10-spc* regions contained rpl23-rpl2-rps19-rpl22-rps3-rpl16-rps17-rpl14-rpl5-rps8[40] which were identical to that in the *E.coli* operons[41], even in prokaryotic genomes [42], this location in cpDNA may come from archaebacteria to the green plant. What is more interested, in *Arabidopsis thaliana, psbB-psbT-psbN-psbH-petB-petD* and *rps3-rpl22-rps19-rps2-rps23* have the same transcriptomes expression pattern which was exactly different from *rpoA-rps11-rpl36-rps8-rpl14-rpl16* under various biological conditions[a1].

Near the *S10–spc*–alpha operon locus, there existed another larger block: *atpA-atpF-atpH-atpI- rps2-petN-psbM-psbD-psbC-psbZ-rps14-ycf3-rps4-ndbJ-ndhK-ndhC-atpE-atpB-rbcL-accD-psaI-ycf4-cemA-petA-psbJ-psbL-psbF-psbE-petL-petG-psaJ-rpl33-rps18-rpl20-psbB-psbT-psbN-psbH-petB-petD-rpoA-rps11-rpl 36* which are ATP synthase, Phytosystem II, Cytochrome and Ribosomal genes.

### Dynamic Evolution of IR in green plant

IR in green algal showed large fluctuation in size from 6.8 kb to 45.5kb, and sustained losses in major groups of green algal but in the green lineage, IR underwent expansion [43, 44]. When compared the IRs regions in green plants, not only for Eudicots but also all angiosperm, chloroplast have experienced an expansion at the end of IRs. For angiosperm, *rps19-rpl2-rpl23-ndhB-rps7-rps12* gene copies were newly acquired in IRs, and there is always a big block connected with IRb. In the fern and horsetails, *ndhB-rps12-rps7* appeared but not in IRs by rearrangement. In Gymnosperm except for Pinales, *ndhB-rps7-rps12* block was copied and inverted to form *rps19-rpl2-rpl23-ndhB-rps7-rps12*, but we cannot tell whether the *ndhB-rps7-ycf12* block was gained by fusion or rearrangement. From Amborellales and Nymphaeales, *rps19-rpl2-rpl23-ndhB-rps7-rps12* gene copies were found in IRs, especially in Nymphaeales which still contains the largest IRs block.

### Congruence and conflict in phylogenetic trees with other studies

There are two previous chloroplast’s phylogenetic analysis of Ruhfel et al. (2014) [28] and Gitzendanner, M.A., et al (2017) [29] where they used 360 and 1879 taxa to study the green plants respectively. Most topologies of our phylogenetic tree were consistent, however, there were some differences between our results and those of the two studies. Based on the AA analysis of Gitzendanner, M.A., et al (2017) recovered Bryophyte clade as monophyletic, which is similar to our results but with an uncertain relationship to Lycophytes and fern. In our matrix AA analysis, we found Bryophyte + Lycophytes are sister to ferns and horsetails (UFboot = 100%). With matrix nt123, hornworts, mosses, and liverworts were identified as successive sister lineages of *Tracheophytes* (UFboot = 100%). Both these two topologies were well supported by previous research [18, 45]. In our study, Magnoliids were placed at the outside of Monocots in matrix nt123 (UFboot = 100%), but nested between monophyletic Monocot and Eudicot in matrix nt12 (UFboot = 59%) and matrix AA (UFboot = 35%). But, from the gene constitution in Magnoliids, they are similar to both Monocot and Eudicot. In Matthew et al. (2017) analysis, Magnoliids and Chloranthales form a weakly supported clade (BS = 61%) that is sister to a clade of Monocots (BS = 100%) and Eudicots. When we combined the dataset from Gitzendanner, M.A., et al (2017) with our AA sequences and re-analyzed, Magnoliids moved outside of the Monocots (SH-alrt =97.5%, UFboot = 95%). Ruhfel et al. (2014) recovered Ceratophyllales as sister to the Monocots using matrix nt12 with low support (BS = 52%) and when using matrix nt123, Ceratophyllales was placed between the Monocots and Eudicots (BS = 52%). We recovered Ceratophyllales + Chloranthales as sister to the Eudicots using nt123 data (UFboot=73%, UFboot=99%), but when using matrix nt12 and AA data, only Ceratophyllales sister to the Eudicots (UFboot=100%).

## Conclusion

The structure of chloroplast genomes is mostly consistent in green plants and formed several gene clusters and gene block except in Chlorophyta. This structural conservatism might be a result of the common genes between cyanobacteria or the same function categories are more likely to form a gene cluster. Topologies of phylogenetic tree of green plants, more extensive taxon indeed increased the phylogenetic resolution for some controversial clades. Matrix nt12 data produced similar results with matrix AA data, matrix nt123 data affected the position of Bryophyte and Magnoliids. In general, for some controversial clades that are deep within green plants, such as, Bryophyte, dense taxon sampling did not improve phylogenetic accuracy anymore, data set will effort the topologies. Thus, resolving the controversial deep-level clades, simply increasing taxon sampling may not be necessary. In addition, chloroplast genome analysis alone seems unlikely to solve the relationship of these controversial clades. Using large numbers of nuclear genes or selecting the nuclear genes with stronger phylogenetic signals may help to solve these deep-level questions.

## Methods

### Taxon and Gene Sampling

The complete or nearly complete chloroplast genomes of 3246 species in GenBank (as of May 18, 2018) and 731 species of Ruili Botanical Garden were retrieved and used as the raw data (Liu et al 2019). The duplicated samples and the species with incorrect annotation or significant gene losses were excluded from the analysis. Six problematic species (*Monoraphidium neglectum*, CM002678; *Nothoceros aenigmaticus*, NC-020259; *Nymphaea ampla*, NC-035680; *Allium sativum*, NC-031829; *Bambusa oldhamii*, NC-012927 and *Potentilla micrantha*, HG931056) were subjected to re-annotation with GeneWise v2.4.1 (Birney et al. 2000). In addition, 50% missing genes of a species and 50% missing species of a gene was also set as the filter elements, and a total of 3654 species with 72 conserved protein-coding genes were obtained for the further analysis. The 298 families from 111 orders of Chlorophyta, Charophyta, Bryophytes, Pteridophyta, Bryophytes, Gymnosperm and Angiosperms were used to represent most major lineages of green plants, and 6 species of Rhodophyta as outgroups were used. The details (species name, family names, order names, genome size, and accession numbers) of 3654 chloroplast genomes are listed in Supplementary Table S1.

### Sequence Alignment

DNA sequences of 72 protein-coding genes were extracted from each genome according to the annotation files. A total of three data sets were made, which included the matrix of all nucleotide positions (nt123), the matrix containing only the first and second codon positions (nt12), and the matrix of amino acids (AA). The nucleotide sequence of each coding gene was processed individually with TranslatorX using MAFFT v7.017 (Katoh et al. 2013) to align the aa sequences, and then the TrimAL software was used to trim the poorly aligned positions with the gappyout option. The genes with 50% missing data were also discarded on the basis of the sequence alignment for further filtration, and the final alignments of each gene were obtained by repeating the aforementioned steps. The nucleotide and amino acid sequence alignments of 72 protein-coding genes were connected subsequently, and a total of 44,187 sites of the ntAll matrix, 29,458 sites of nt12 matrix and 14,724 amino acid positions of AA matrix were used for further analysis.

### Phylogenetic analyses

Three datasets containing 72 protein-coding genes of 3654 species with no partitioning strategies were used to reconstruct the phylogenetic tree based on IQ-TREE with 5000 ultrafast bootstrap replicates and 1000 bootstrap replicates for SH-aLRT, together with GTR+F+R10 model for nucleotide sequences and JTT+F+R10 model for amino acids sequences. ML analysis was also conducted using RAxML v8.2.4 (Stamatakis 2014) under the GTRCAT model for nucleotide and PROTGAMMAWAG model for amino acids. The 100 bootstrap replicates were set to test the reliability of each node for ML analysis.

The optimal partitioning scheme was referred according to Brad et. (2014. For the AA data, we partitioned the data sets by gene (72 partitions). For the nucleotide (nt123), we used program PartitionFinder (ref) to find the best partitioning strategies by each codon position within each gene, to reduce the computational burden, only 731 species of Ruili Botanical Garden were used for PartitionFinder. The 148 partitions were selected with lnL: -3013791.4635925293, AICc: 6034073.71459. Partitioning strategies for both AA data and nt123 to construct phylogenetic tree were unable to complete due to time limitations resulting from the large samples of our data.

To verify the topologies of the phylogenetic tree, the amino acids sequences of 72 genes of 1901 samples in former research (Gitzendanner et al. 2018) were downloaded to analyze along with our data using the IQ-TREE. The Tree-doctor was used to obtain the simplified trees in order levels. The species of Rhodophyta was set as outgroups to re-root the result, and the iTOL (https://itol.embl.de/) was used for data visualization.

### Evolutionary Rate Estimation

According to the gene losses of ndh subunits, 37 species in all 14 clades (10 of 37 species lost all ndh genes) were selected as the representative species, and the sequence of 14 photosynthesis genes existing in all 37 species was selected. The alignments and the topology derived from IQ-TREE of all nucleotide positions (nt12) of each gene and the gene set in the selected species were used to perform the evolutionary rate analysis. The codeml of PAML was used to estimate the ratio of nonsynonymous to synonymous nucleotide substitutions (dN/dS) for each branch of 14 genes corresponding to photosynthesis.

## Data availability

The data that support the findings of this study have been deposited in the CNSA (https://db.cngb.org/cnsa/) of CNGBdb with accession code CNPhis0000538.

## Acknowledgements

This work was supported by the grants of Basic Research Program, Shenzhen Municipal Government of China (No. JCYJ20160331150739027 and No. JCYJ20150831201123287), and the Guangdong Provincial Key Laboratory of Genome Read and Write (Grant No. 2017B030301011).

## Table legends

Table 1. Summary of consensus topologies derived from the three data sets.

## Supporting information

**Table S1.** The gene blocks frequency in Streptophyta

**Table S2.** The detailed information and characters of the species used in this study.

**Table S3.** The gene blocks frequency in Streptophyta

**Table S4.** The hexadecimal colors used in this study.

**Table S5.** The 37 selected species for the analysis of dN/dS.

**Fig S1.**
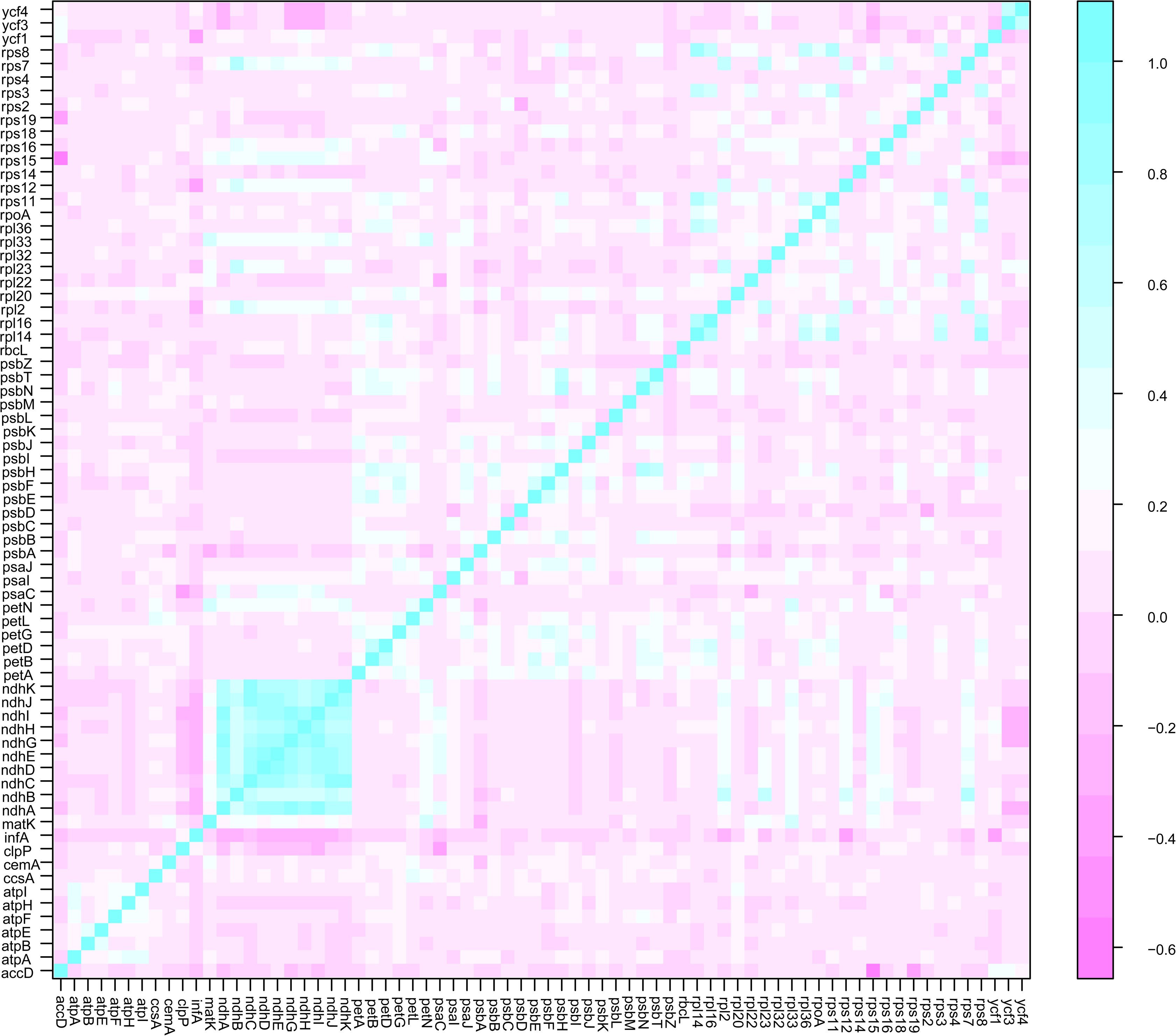
Gene content in the green plants, the correlation of the genes was calculated by the Spearman method.

**Fig S2.**
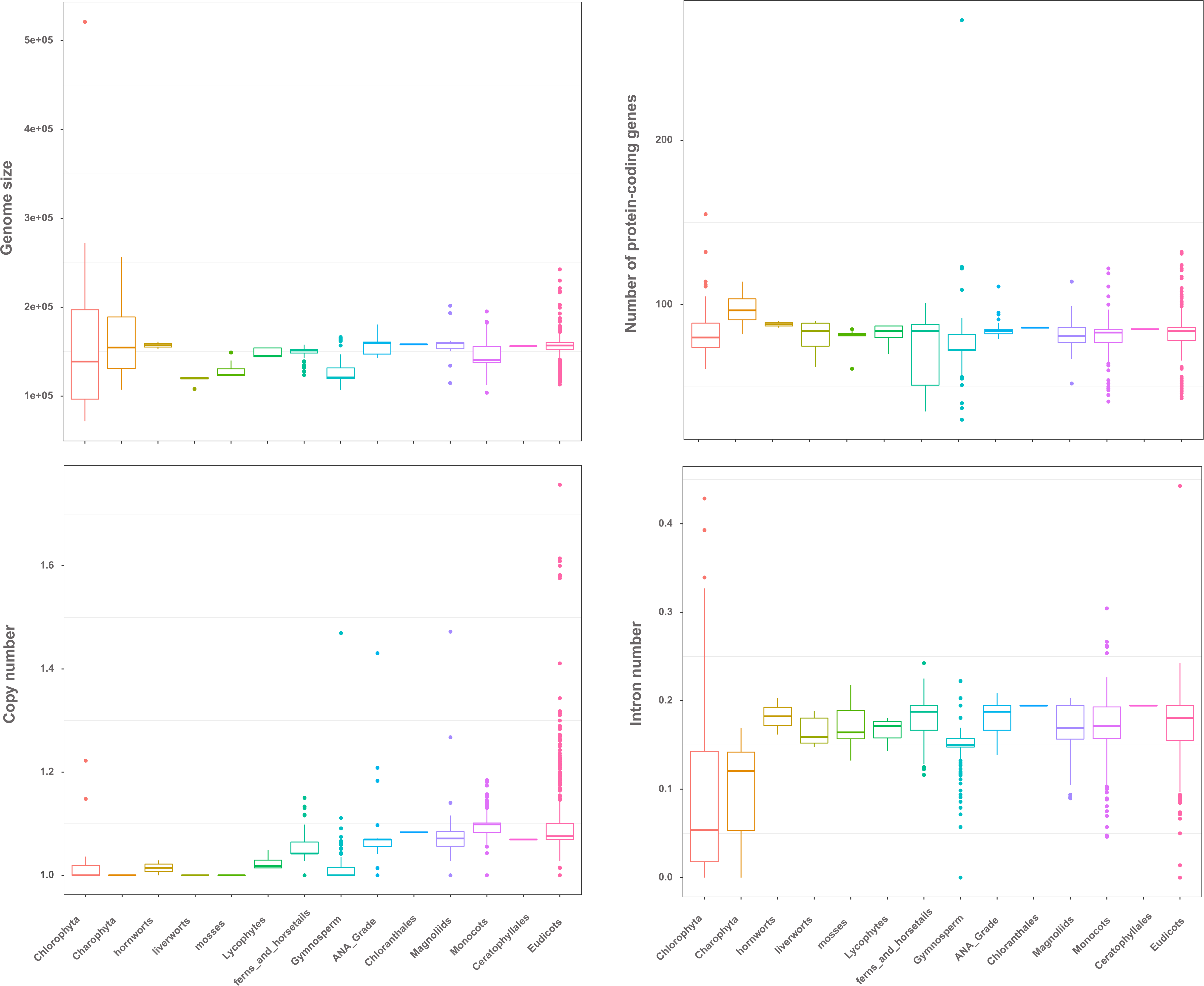
Characters of the chloroplast genome in the green plant. The box plots represent the median (central line), first and third quartiles (box), and outliers (dot).

**Fig S3.**
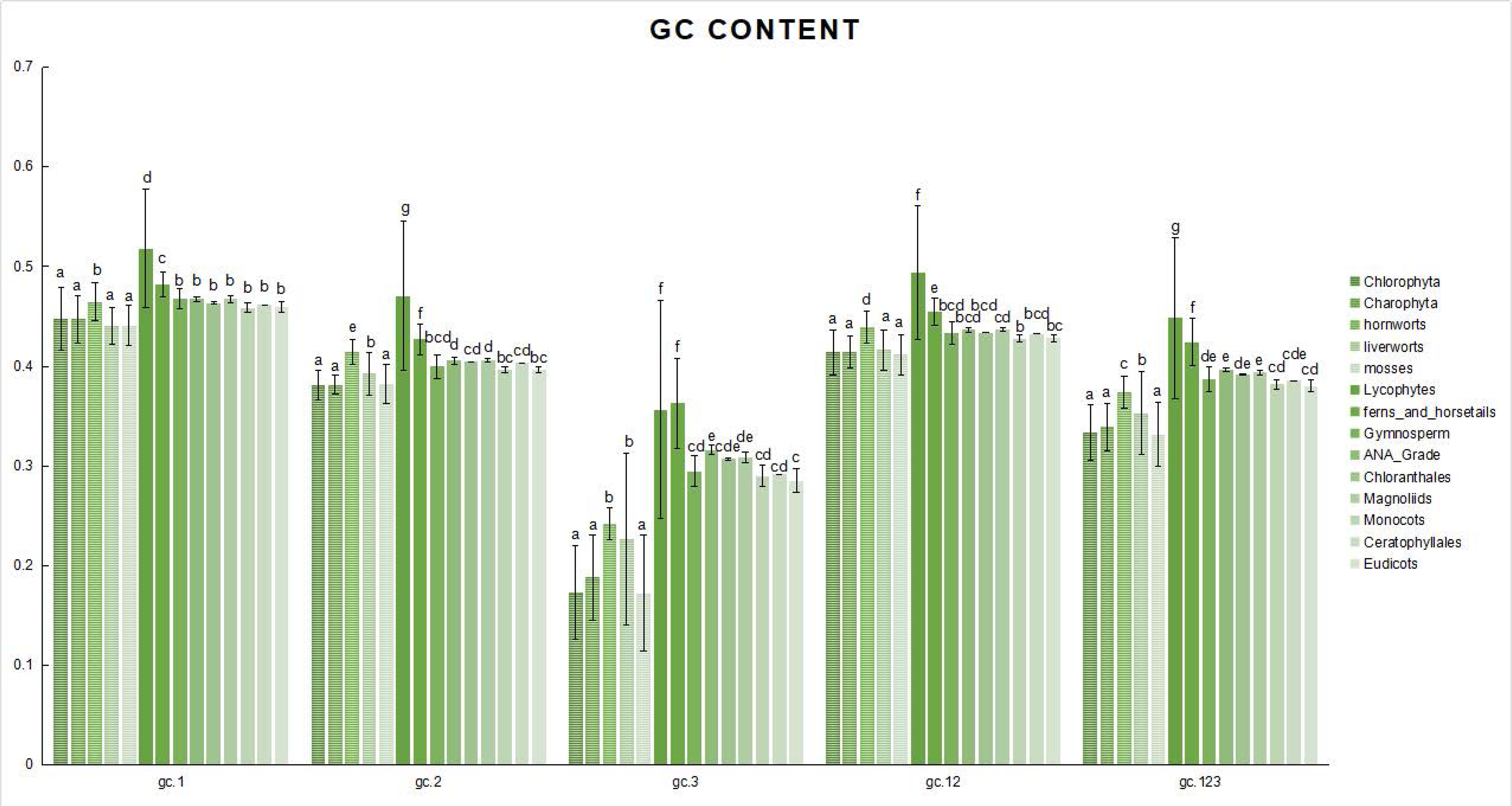
GC content of different codon positions in green plant. The One-Way ANOVA and other statistical analyses were performed in SPSS software, and the lowercase represents the significance between 14 clades (p < 0.01).

**Fig S4.**
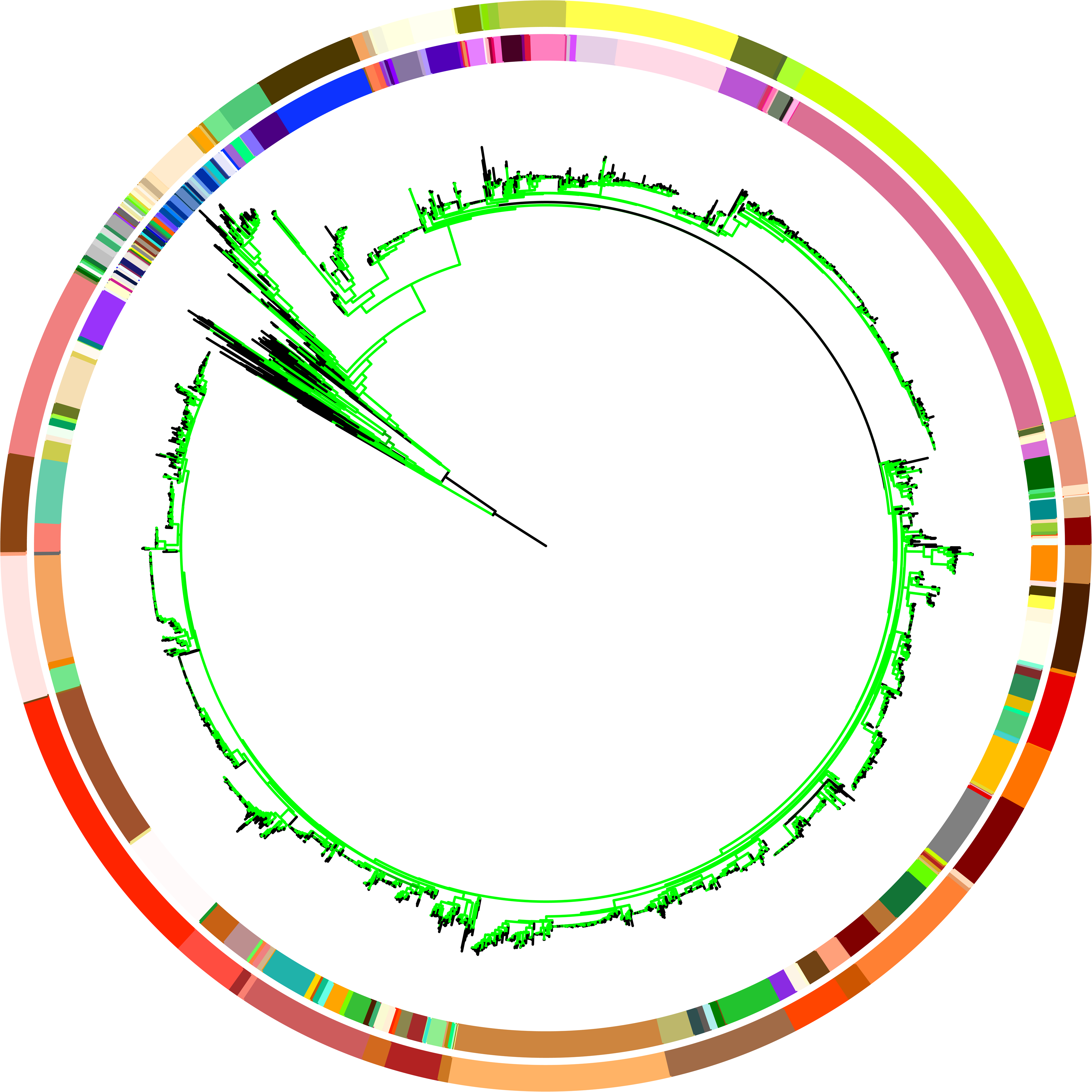
Chloroplast phylogenomic tree based on the matrix nt123 of 72 protein-coding genes of 3654 green plants and six Rhodophyta using IQTREE. The colors on the internal circle indicate different families, while the colors on the external circle indicate different orders.

**Fig S5.**
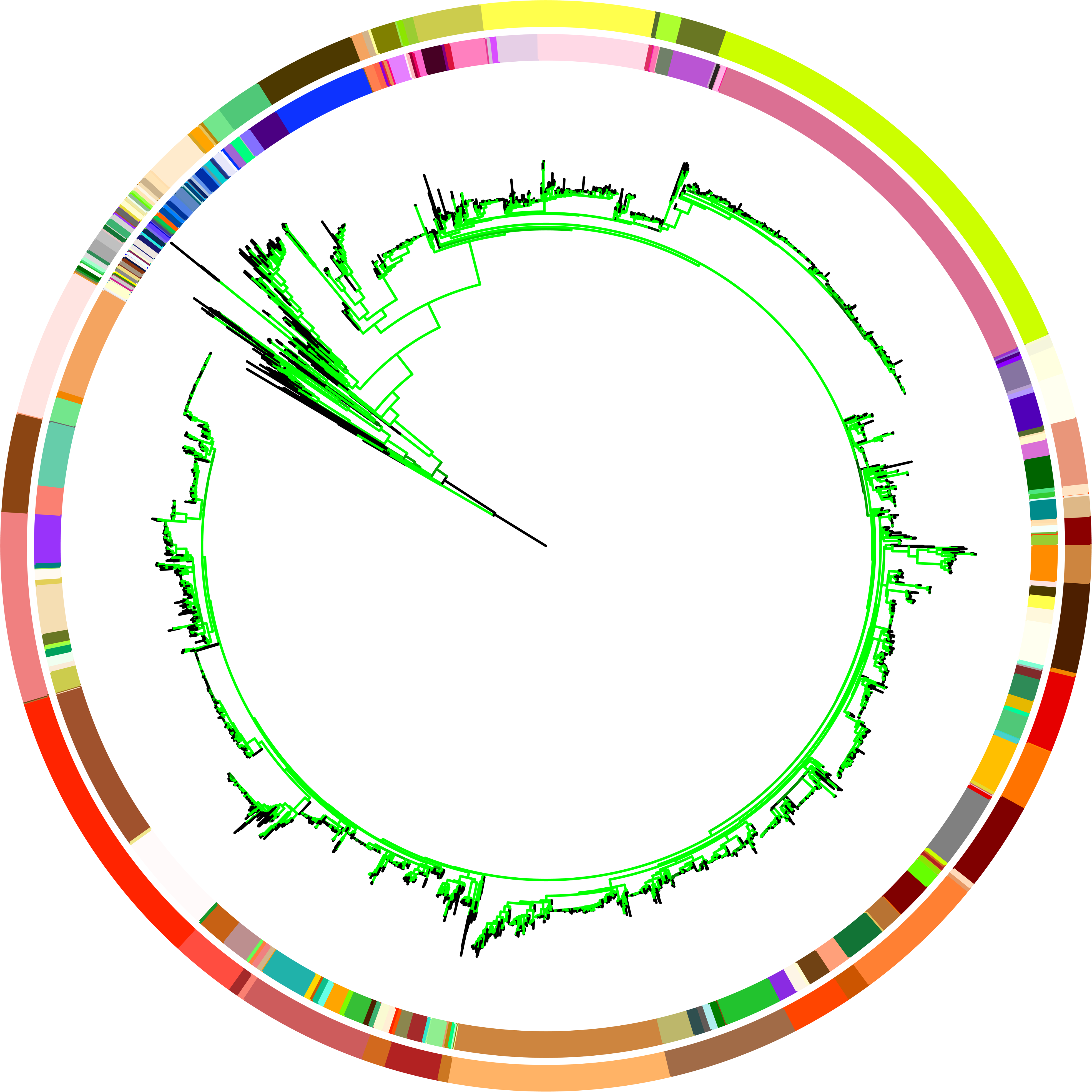
Chloroplast phylogenomic tree based on the matrix aa of 72 protein-coding genes of 3654 green plants and six Rhodophyta using IQTREE. The colors on the internal circle indicate different families while the colors on the external circle indicate different orders.

**Fig S6.**
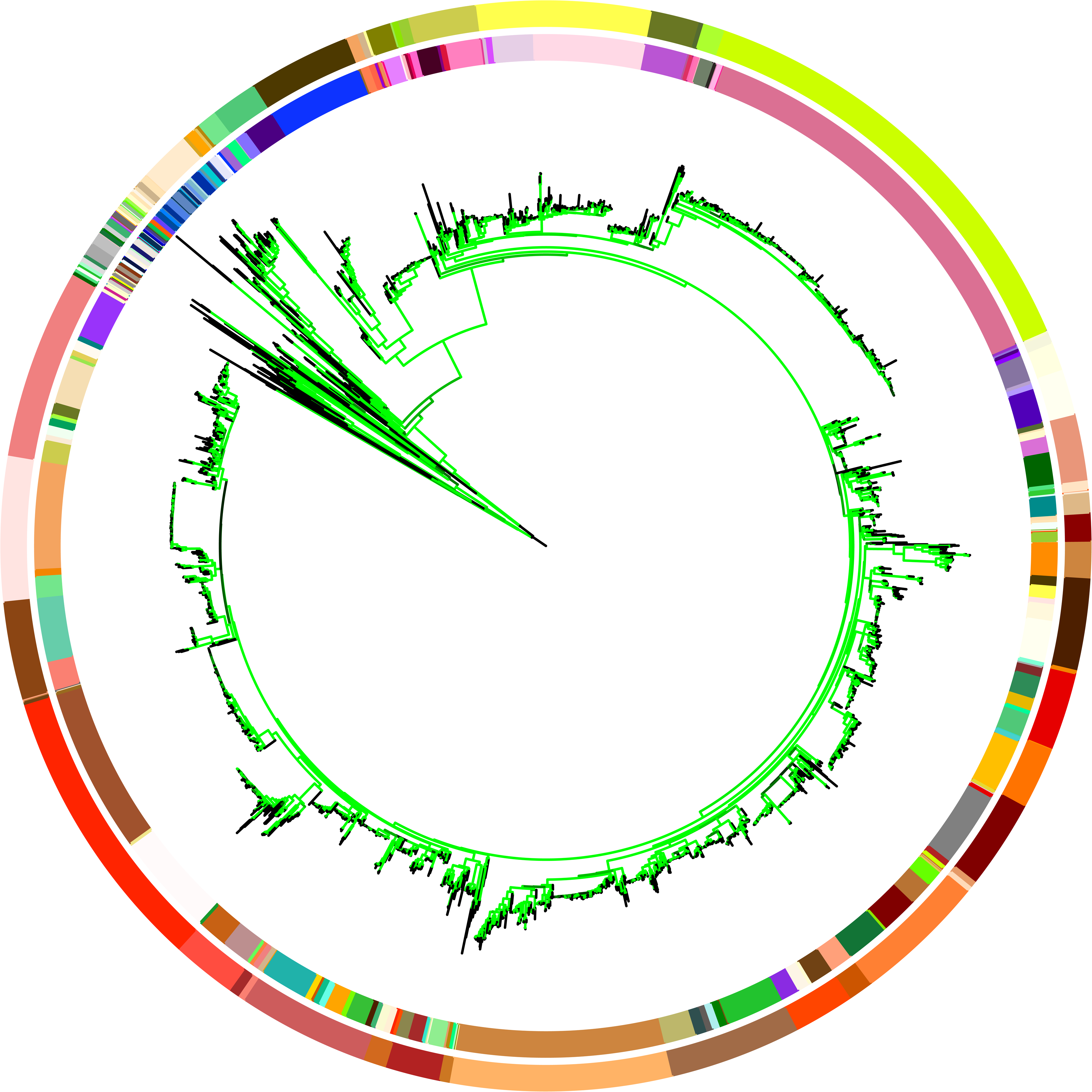
Chloroplast phylogenomic tree based on the matrix nt12 of 72 protein-coding genes of 3654 green plants using RaXML. The colors on the internal circle indicate different families while the colors on the external circle indicate different orders.

**Fig S7.**
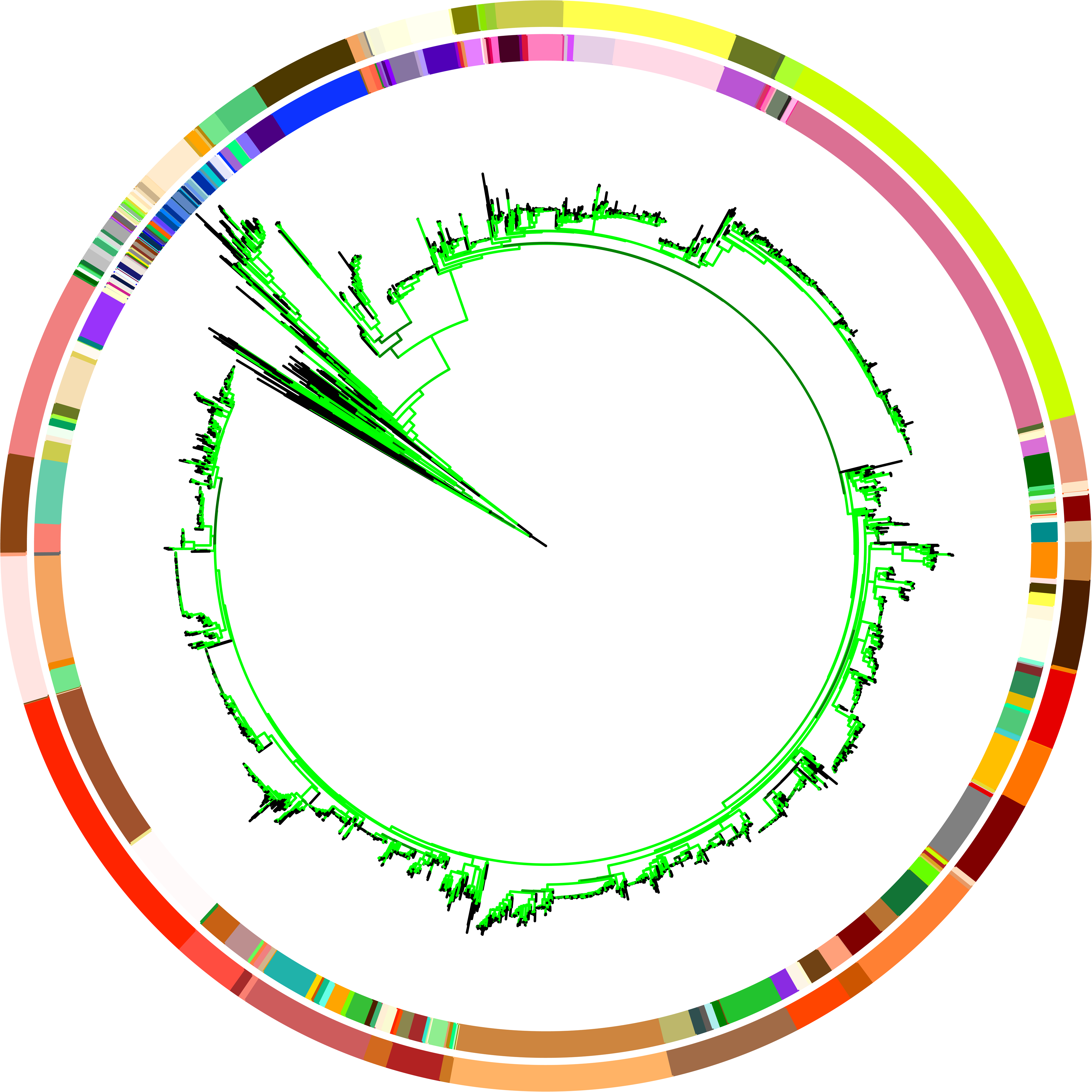
Chloroplast phylogenomic tree based on the matrix nt123 of 72 protein-coding genes of 3654 green plants using RaXML. The colors in the internal circle indicate different families while the colors in the external circle indicate different orders.

**Fig S8.**
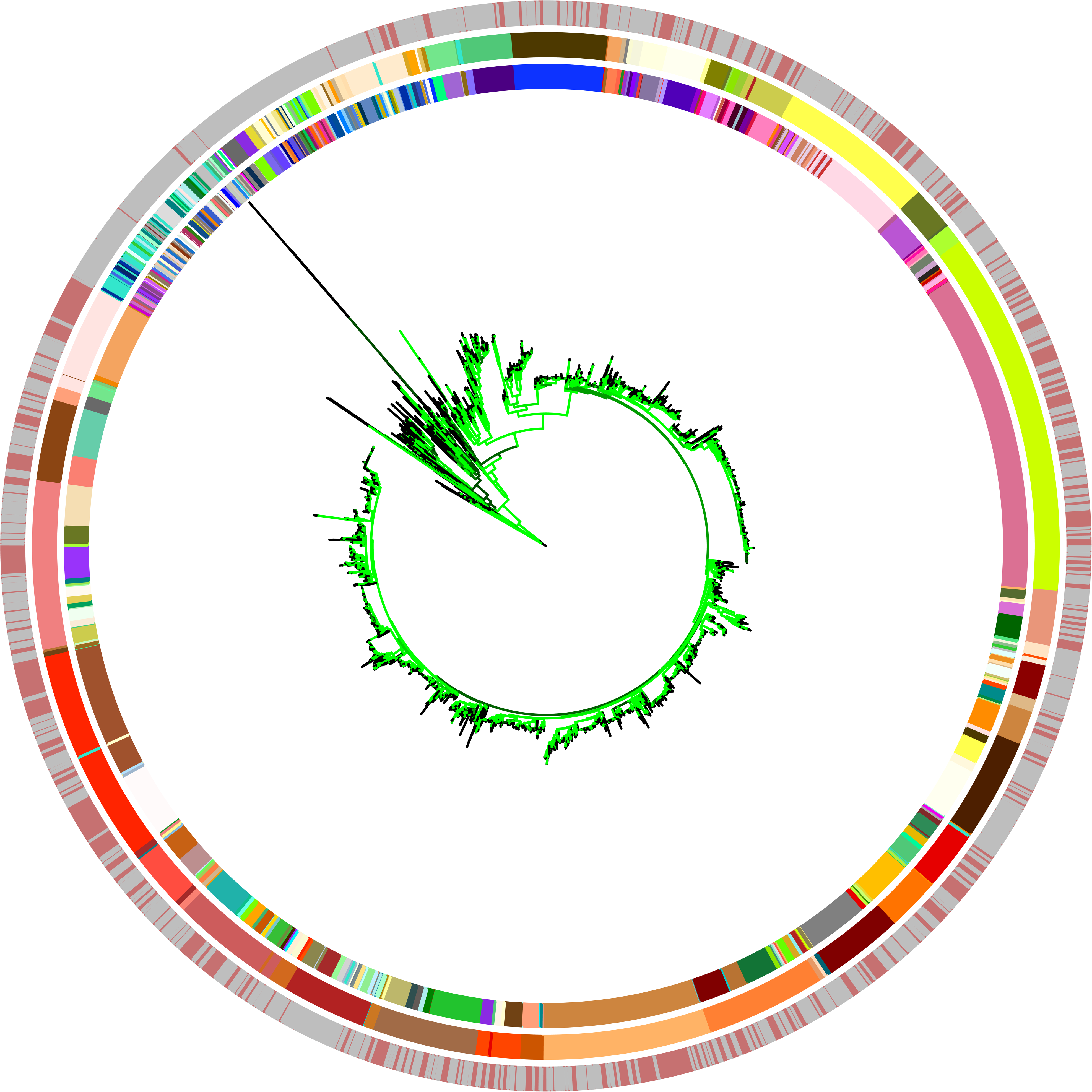
Chloroplast phylogenomic tree based on the matrix aa of 72 protein-coding genes of 3654 green plants and 1901 species in the former research using IQTREE. The colors on the internal circle indicate different families while the colors on the external circle indicate different orders.

**Fig S9.**
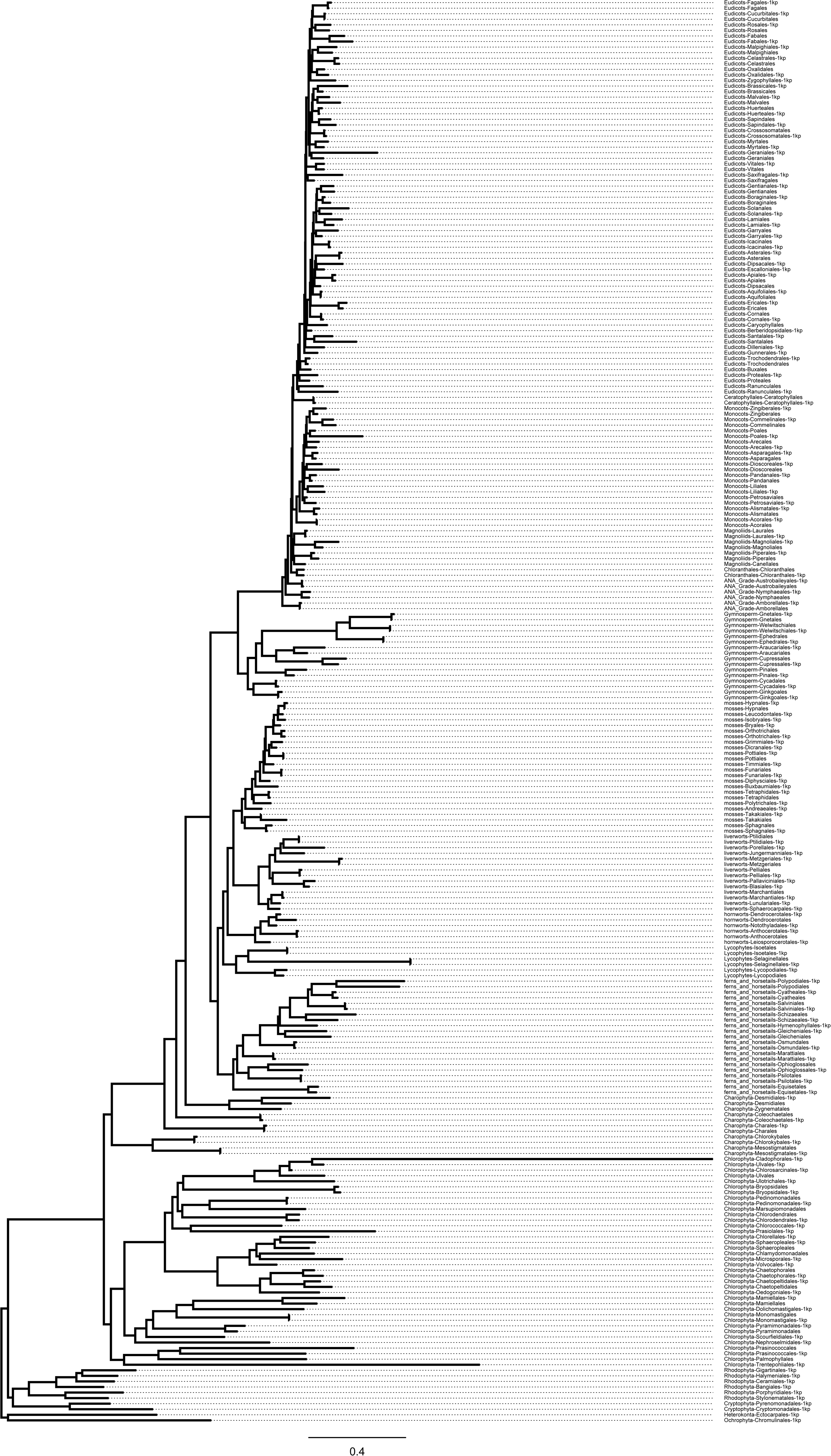
Summary of the phylogenomic tree based on matrix aa of 72 protein-coding genes of 3654 green plants and 1901 species obtained from earlier reports using IQTREE.

**Fig S10.**
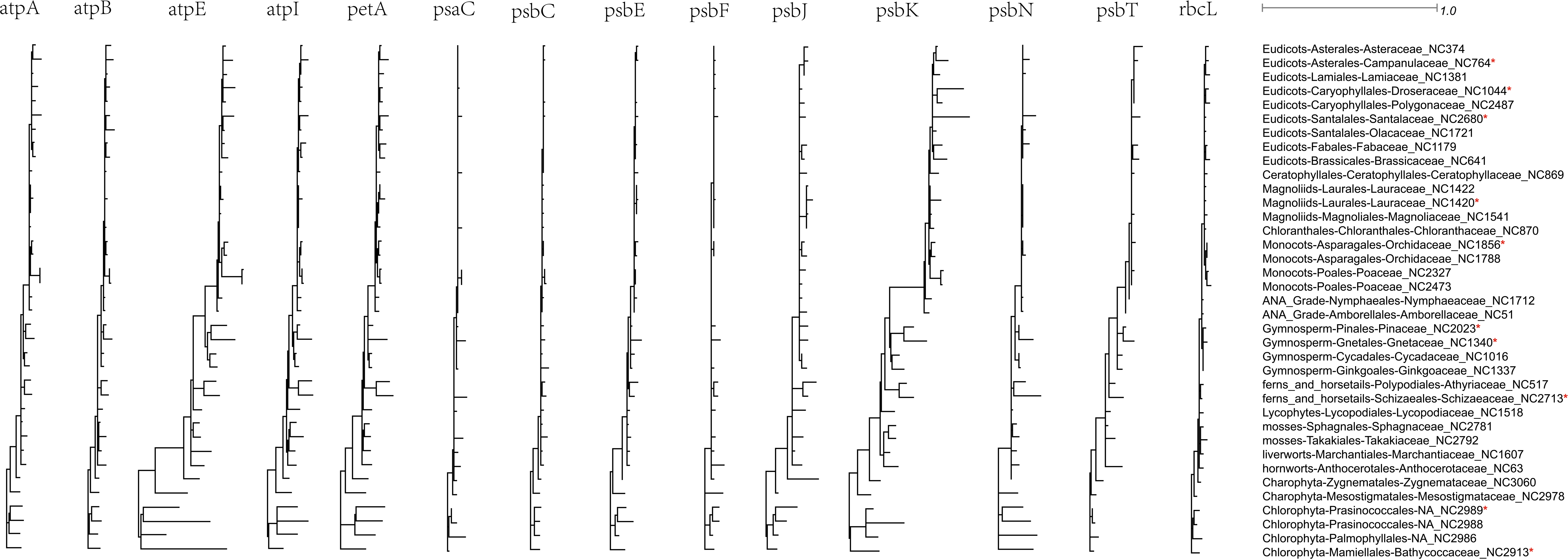
Phylogenomic tree of dN for 14 photosynthesis genes in 37 species. “*” indicated the species without ndh genes.

**Fig S11.**
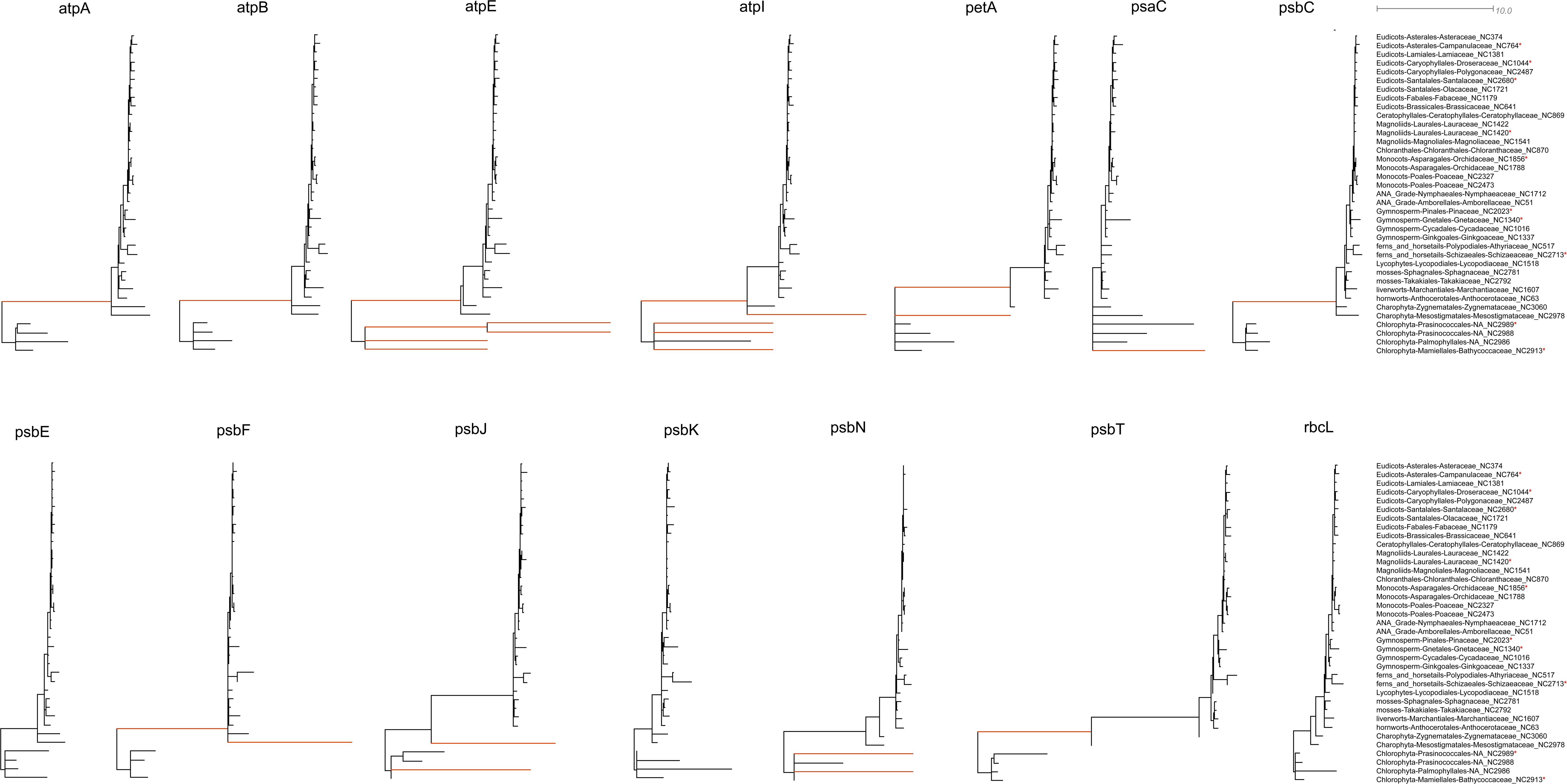
Phylogenomic tree of dS for 14 photosynthesis genes in 37 species. The red branch stands for the length of the branch which was changed to 10. “*” indicated the species without ndh genes.

